# B7-H1(PD-L1) confers chemoresistance through ERK and p38 MAPK pathway in tumor cells

**DOI:** 10.1101/308601

**Authors:** Xiaosheng Wu, Yanli Li, Xin Liu, Siyu Cao, Susan M. Harrington, Chunhua Chen, Aaron S. Mansfiled, Roxana S. Dronca, Sean S. Park, Yiyi Yan, Eugene D. Kwon, Liewei Wang, Kun Ling, Haidong Dong

## Abstract

Development of resistance to chemotherapy and immunotherapy is a major obstacle in extending the survival of patients with cancer. Although several molecular mechanisms have been identified that can contribute to chemoresistance, the role of immune checkpoint molecules in tumor chemoresistance remains underestimated. It has been recently observed that overexpression of B7-H1(PD-L1) confers chemoresistance in human cancers, however the underlying mechanisms are unclear. Here we show that the development of chemoresistance depends on the increased activation of ERK pathway in tumor cells overexpressing B7-H1. Conversely, B7-H1 deficiency renders tumor cells susceptible to chemotherapy in a cell-context dependent manner through activation of the p38 MAPK pathway. B7-H1 in tumor cells associates with the catalytic subunit of a DNA-dependent serine / threonine protein kinase (DNA-PKcs). DNA-PKcs is required for the activation of ERK or p38 MAPK in tumors expressing B7-H1, but not in B7-H1 negative or B7-H1 deficient tumors. Ligation of B7-H1 by anti-B7-H1 monoclonal antibody (H1A) increased the sensitivity of human triple negative breast tumor cells to cisplatin therapy in vivo. Our results suggest that B7-H1(PD-L1) expression in cancer cells modifies their chemosensitivity towards certain drugs and targeting B7-H1 intracellular signaling pathway is a new way to overcome cancer chemoresistance.

## INTRODUCTION

Development of resistance to chemotherapy and immunotherapy is a major obstacle in prolonging the survival of patients with cancer. Although several molecular mechanisms have been identified to contribute to chemoresistance (1, 2), the role of immune checkpoint molecules in the development of tumor chemoresistance has not been fully recognized. Traditionally, the emergence of chemoresistance and immunoresistance are considered as parallel and unrelated events. However, recent studies indicate that the overexpression of some immune checkpoint molecules (e.g. B7-H1/PD-L1 or B7-H3) not only negatively influences antitumor immunity, but also renders tumor cells resistant to chemotherapeutic agents(3–5). Elevated expression of B7-H1(PD-L1) is predictive of an aggressive disease course, including increases in rate of disease progression and cancer-related mortality in a variety of cancers (ovary, breast, liver, pancreas, esophagus, colon and lung)(6, 7). Although the poor prognosis of B7-H1 positive tumors has been attributed to B7-H1’s immune-suppressive function, the emerging role of B7-H1 in tumor chemoresistance may provide important clues in combating cancer.

It has been previously observed that the overexpression of B7-H1 confers chemoresistance in human multiple myeloma and breast cancers(3, 4) while B7-H1 knockdown in human lymphoma and breast tumor cell lines renders them sensitive to chemotherapeutic drugs(4, 8). These results suggest that B7-H1-mediated chemoresistance may be cell-context dependent. However, the mechanism by which B7-H1 governs tumor chemoresistance remains unclear. Identification of this mechanism would enable us to identify new therapeutic targets to combat chemoresistance. In this study, we examined B7-H1 expression in several human and mouse tumor cell lines in context of their drug sensitivity in vitro and in vivo. We reported that overexpression of B7-H1 led to an increased activation of ERK and survivin in tumor cells through an association with DNA-PKcs, and B7-H1 deficiency renders tumor cells susceptible to chemotherapy in a cell-context dependent manner through activation of the p38 MAPK pathway and Bcl-2. Our results provided new insights into the B7-H1-mediated cancer chemoresistance and identified a potential therapeutic target to overcome chemoresistance in human cancer.

## RESULTS

### B7-H1 overexpression renders tumor cell resistant to chemotherapy

To explore how B7-H1 affects sensitivity to chemotherapy in cancer cells, we transfected a B7-H1 negative human melanoma cell line (624mel) with a full-length human B7-H1 cDNA construct (9). The B7-H1 cDNA transfected and mock-transfected 624mel cells were confirmed by confocal microscopy (Fig. 1A) and flow cytometry (SF1. A) for their B7-H1 expression status, respectively. B7-H1 protein was found to be located both on the cell membrane and in the cytoplasm of the tumor cells, whereas neither PD-1 nor CD80, two receptors of B7-H1, was detected in 624mel cells (SF. 1A). The transfection of B7-H1 did not affect the proliferation of 624mel cells as shown by their comparable Ki67 expression and cell growth pattern (SF1. B-C).

**Figure 1.**
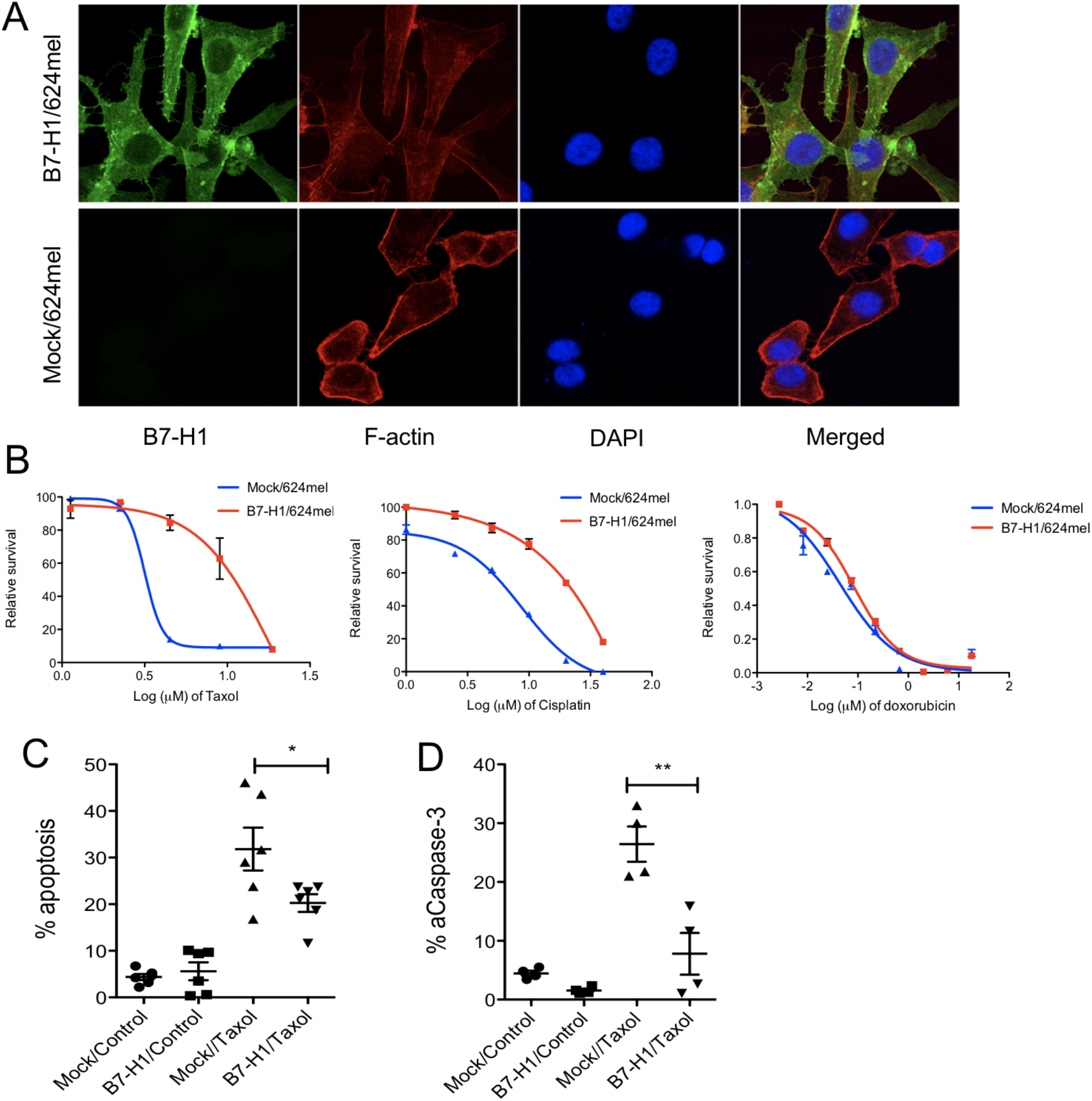
B7-H1 overexpression conferred drug resistance on tumor cells. (A) B7-H1 expression by B7-H1 transfected 624mel tumor cells. (B) Relative survival was determined by MTS assay. The p-value for area under the curve (dose-response curves) is significant (p<0.01) in treatment with taxol (paclitaxel) and cisplatin, no significant difference in treatment with doxorubicin. One of three experiments is shown. (C-D) Apoptosis of tumor cells treated with taxol (2 μg/ ml, 48 hours) was analyzed by TMRE and Annexin V staining (C) and intracellular staining for active caspase-3 (D). Numbers are percentages of gated population. *P<0.05, **P<0.01 compared between Mock and B7-H1/624mel cells.

Despite prior reports indicating that overexpression of B7-H1 renders tumor cells resistant to chemotherapeutic agents (3, 4), it is not clear whether elevated B7-H1 expression specifically enhances the resistance toward certain type(s) of drugs. To this end, we tested the sensitivities of our B7-H1 overexpressing tumor cells to various chemotherapeutic drugs including paclitaxel (Taxol, targeting microtubules and blocks mitosis), cisplatin (crosslinks DNA to stop replication of DNA) and doxorubicin (topoisomerase II inhibitor and generation of free radicals) in cytotoxicity assays. Specifically, identical numbers of 624mel cells transfected with B7-H1 or the control vector were incubated with increasing concentrations of each of these chemotherapy drugs (Fig. 1B). After 72 hours of culture, the viabilities of these cells were measured using an MTS assay(10). Our results show that the 624mel cells expressing B7-H1 were more resistant to paclitaxel and cisplatin comparing to the control cells; but exhibited similar sensitivity to doxorubicin as the control cells (Fig 1. B). Since a mechanism of action of most cytotoxic drugs is to induce apoptosis (11, 12), we measured and compared the apoptosis of B7-H1/624mel and Mock/ 624mel tumor cells after treatment with paclitaxel. Apoptosis was measured by the binding of Annexin V (AV) and the levels of tetramethylrhodamine ethyl ester (TMRE, a marker for mitochondria membrane potential)(13). B7-H1/624mel cells had 1.5 fold less apoptotic cells compared to Mock/ 624mel cells following the treatment with paclitaxel (Fig. 1C). The level of active caspase-3 (an key molecule of apoptosis) also was lower in B7-H1/624mel cells compared with Mock/ 624mel cells (Fig. 1D). Our results suggest that B7-H1 overexpression renders tumor cells resistant to certain chemo drugs by reducing their pro-apoptotic potentials.

### B7-H1 deficiency renders tumor cell susceptible to chemotherapy in a cell-context dependent way

To examine whether depletion of endogenous B7-H1 would affect the sensitivity to chemotherapy in tumor cells, we took the B7-H1 positive human breast cancer cell line MDA-MB-231 and produced a B7-H1 biallelic knockout subclone by introducing CRISPR/ Cas9 constructs carrying a guide RNA (gRNA) sequence specific to human B7-H1 exon 3. The B7-H1 deficiency in the knockout cells was confirmed by genotyping, confocal microscope, flow cytometry and Western blotting (Fig. 2A-C). We also confirmed that MDA-MB-231 cells did not express B7-H1 receptors PD-1 and CD80 (Fig. 2C). We then tested the sensitivity of B7-H1 deficient MDA-MB-231 cells to various chemotherapeutic drugs, our results showed that the depletion of B7-H1 promoted the sensitivity of the breast cancer cells to the treatments of cisplatin (Fig. 2D). Cisplatin induced more apoptosis and activated caspase-3 in B7-H1 deficient MDA-MB-231 cells than in their wild type counterparts (Fig. 2E-F). Using a dose of cisplatin that did not suppress the growth of WT MDA-MB-231 cells in vivo, we found the same dose of cisplatin significantly suppressed the growth of B7-H1 KO MDA-MB-231 cells (Fig. 2 G-H). Of note, B7-H1 WT and B7-H1 KO MDA-MB-231 tumors had similar growth in vivo in immune deficient mice (Fig. 2 I). We also found anti-B7-H1 antibody (clone H1A) significantly suppressed the growth of B7-H1 positive WT MDA-MB-231 cells but not B7-H1 KO MDA-MB-231 cells, suggesting a B7-H1 specific effect of anti-B7-H1 antibody (Fig. 2 G-H). Similar to B7-H1 deficiency, anti-B7-H1 antibody rendered B7-H1 WT MDA-MB-231 cells more sensitive to cisplatin in suppression of tumor growth (Fig. 2 G-H). In addition to breast cancer cells, we produced B7-H1 knockout human kidney cancer 786-0 and human lung cancer A549 that expressed high and low levels of endogenous B7-H1, respectively (Supplemental Fig. 2). We found that B7-H1 depletion increased susceptibility to cisplatin in 786-0 cells but not in A549 cells (Fig. 2G-H), suggesting that B7-H1 modulates chemosensitivity is universal, but the drug specificity is dependent on tumor cell-context.

**Figure 2.**
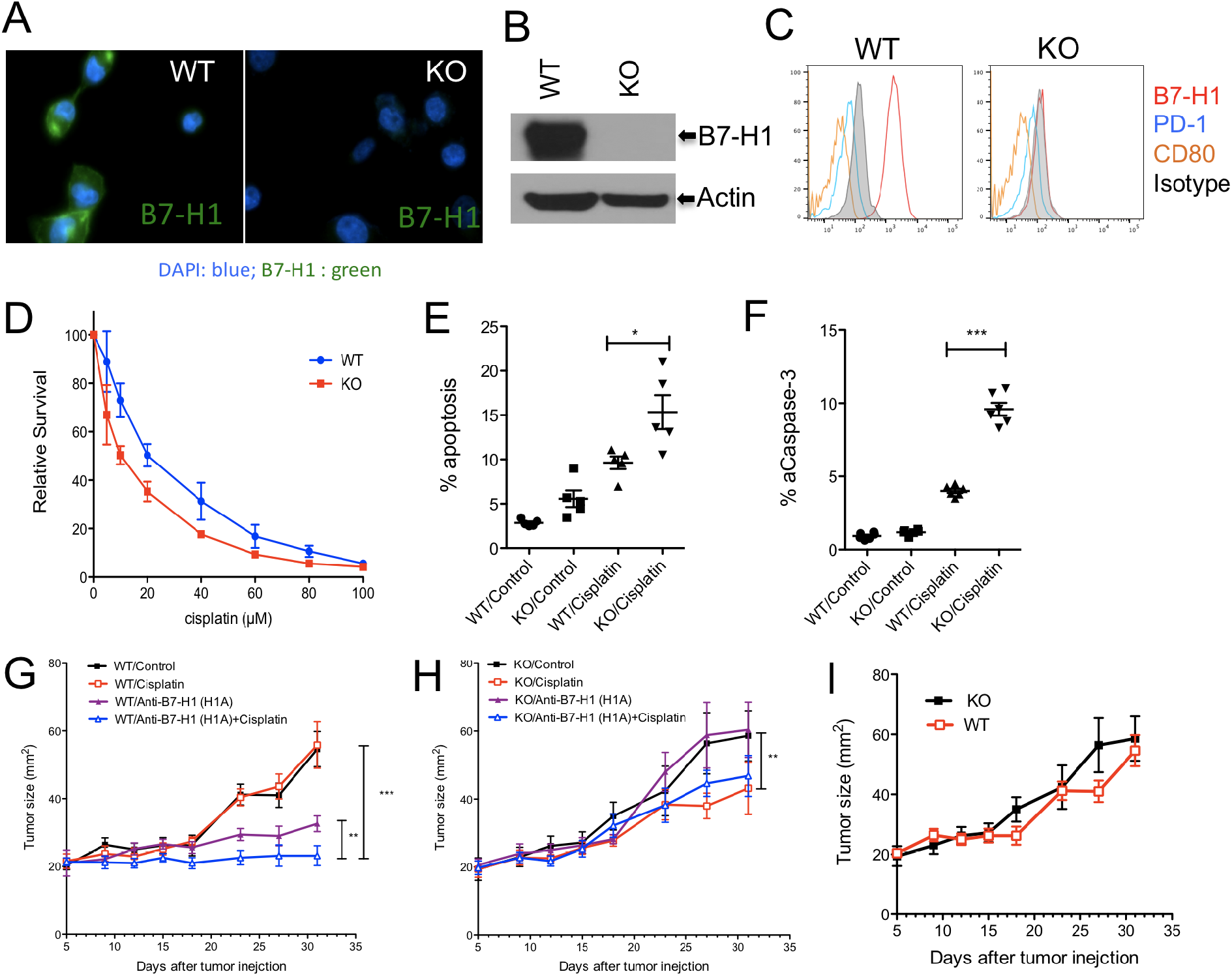
B7-H1 deficiency and B7-H1 antibody made tumor cells sensitive to chemotherapy. B7-H1 deficiency in MBA-MD-231 breast cancer cells was generated by CRISPR/Cas9 and confirmed by confocal microscopy (A), Western blotting (B) and flow cytometry (C). The absence of PD-1 and CD80 expression was confirmed in both wild type (WT) and B7-H1 KO tumor cells (C). (D) B7-H1 deficient MBA-MD-231 cells are more sensitive to cisplatin than WT cells as determined by MTS assay. The p-value for area under the curve (dose-response curves) is significant (p<0.01) in treatment with cisplatin. One of three experiments is shown. (E-F) Apoptosis of tumor cells treated with cisplatin (20 μM, 48 hr.) was analyzed by TMRE and Annexin V staining (E) and intracellular staining for active caspase-3 (F). B7-H1 WT (G) and B7-H1 KO MDA-MB-231 tumor (H) were treated with anti-B7-H1 antibody (H1A) or Cisplatin, or both, in vivo. (I) B7-H1 WT and B7-H1 KO MDA-MB-231 tumors had comparable growth in immune deficient mice without any treatment. *P<0.05, **P<0.01, ***P<0.001 as determined by unpaired t test (in vitro) or two-way ANOVA.

To further determine the cell-specific role of B7-H1 in rendering chemoresistance, we produced B7-H1 deficient mouse breast tumor cells lines from B7-H1 KO mice that spontaneously produce Her2 positive breast cancer. To produce this animal model, we crossed B7-H1 null mice with NeuT Balb/c mice that spontaneously generate Her2 positive breast cancers (Fig. 3A). From the grown out breast cancers, we established Her2 positive breast tumor cell lines with (WT / 5C) or without B7-H1 expression (KO/31A) (Fig. 3B). The presence or absence of B7-H1 expression was confirmed by flow cytometry staining in the established B7-H1 WT and KO breast tumor cell lines (Fig. 3C). B7-H1 deficient tumor cells are more susceptible to the treatment of paclitaxel, but not cisplatin, than B7-H1 WT tumor cells in vitro and in vivo (Fig. 3C-E). Collectively, our studies suggest that B7-H1 renders tumor resistant to chemotherapy in a context of cell type and drug selection.

**Figure 3.**
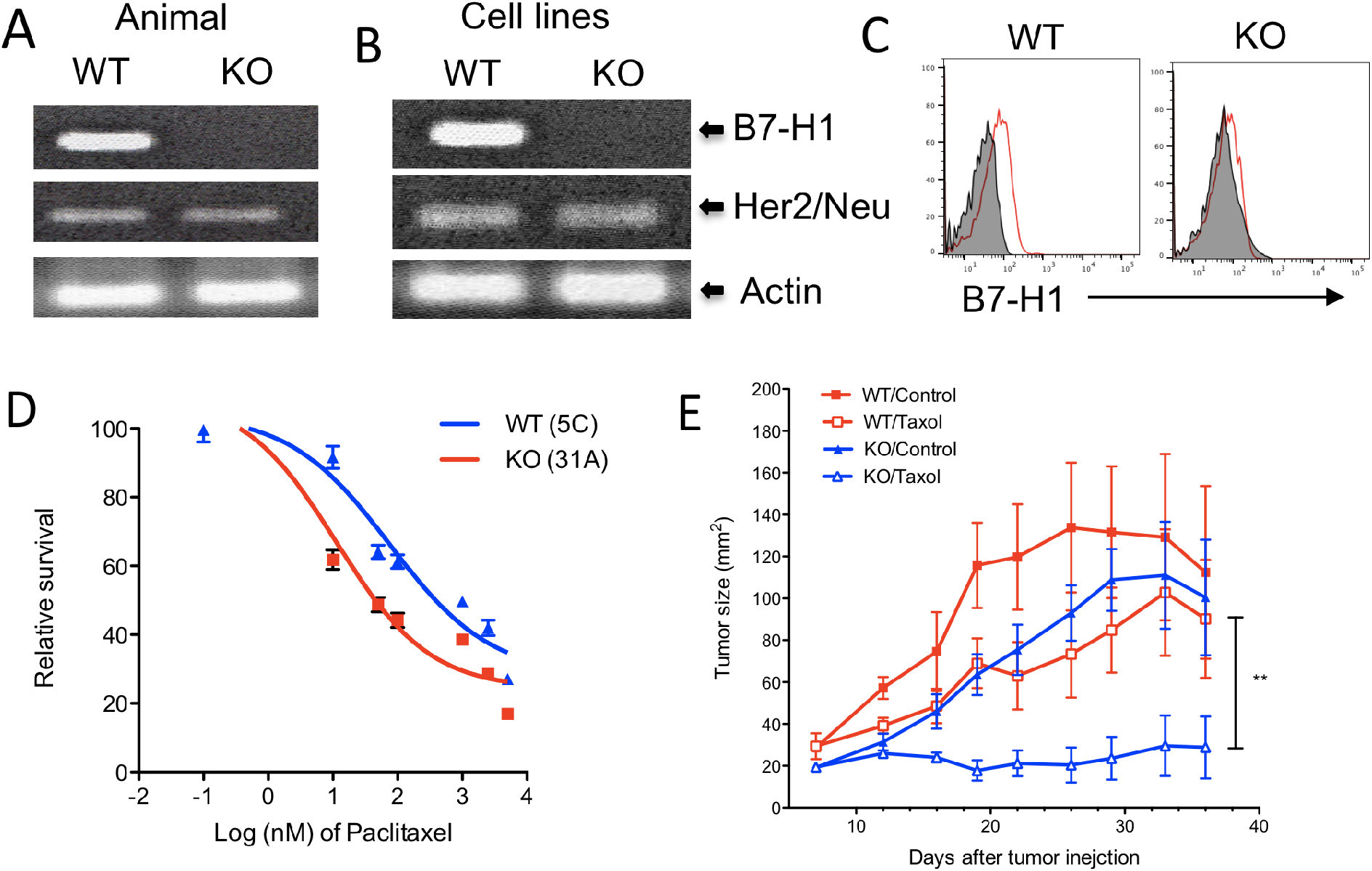
B7-H1 deficient Her2+ breast cancer cells were sensitive to chemotherapy. B7-H1 WT (5C) and B7-H1 KO (31A) mouse breast cancer cell lines were established from wild type WT or B7-H1 KO NeuT mice breast cancer tissues. B7-H1 deficiency and Her2/Neu expression were confirmed by PCR in genomic DNA of mice (A) or breast cancer cell lines (B), and by flow cytometry for cell lines (C). (D-E) B7-H1 KO tumor cells are more sensitive to Taxol (paclitaxel) than WT cells. (E) B7-H1 WT and KO tumor cells were s.c. injected into Balb/c mice at 5×10^5^. On day 8 and 14, paclitaxel (Taxol) was injected i.p. at 25 mg/kg, and control groups were treated with saline. Data show the mean tumor size ± SEM, **P<0.01 compared between WT/Taxol and KO/Taxol (Two-way ANOVA, n=5 group).

### B7-H1 enhances the activation of ERK/MAPK pathway via DNA-PKcs in tumor cells

The absence of B7-H1 receptors PD-1 and CD80 expression in human 624mel and MDA-MB-231 cells excluded the possibility of ligand/ receptor (B7-H1/PD-1 or B7-H1/CD80) mediated cell signaling through PD-1 or CD80 (SF. 1A and Fig. 2C). This prompted us to examine which intracellular pathways B7-H1 itself may directly mediate to regulate the drug sensitivity of these tumor cells. We previously reported that B7-H1 associates with DNA-PKcs in human T cells (14), here we examined whether B7-H1 would be in association with DNA-PKcs in tumor cells. To that end, we performed immunoprecipitation (IP) and Western blotting using anti-B7-H1 or anti-DNA-PKcs antibody on lysates from B7-H1 expressing tumor cells B7-H1/ 624mel and Karpas 299. When the blot was probed with anti-DNA-PKcs, we identified a band of protein (450 kDa) in IP samples immunoprecipitated with anti-B7-H1 antibody but not with isotype control antibody (Fig. 4A). Conversely, when the blot probed with anti-B7-H1 antibody, a B7-H1 band was readily identified in IP samples immunoprecipitated with anti-DNA-PK (Fig. 4A). In addition, we identified the endogenous association of B7-H1 and DNA-PKcs in human B7-H1 positive MDA-MB-231 breast cancer cells (Fig. 4B). Interestingly, a DNA-PKcs inhibitor (NU7026) that inhibits the catalytic activity of DNA-PKcs, abolished the association of DNA-PKcs and B7-H1 in this breast cancer cell line (Fig. 4B), suggesting the binding is dependent on DNA-PKcs activity.

**Figure 4.**
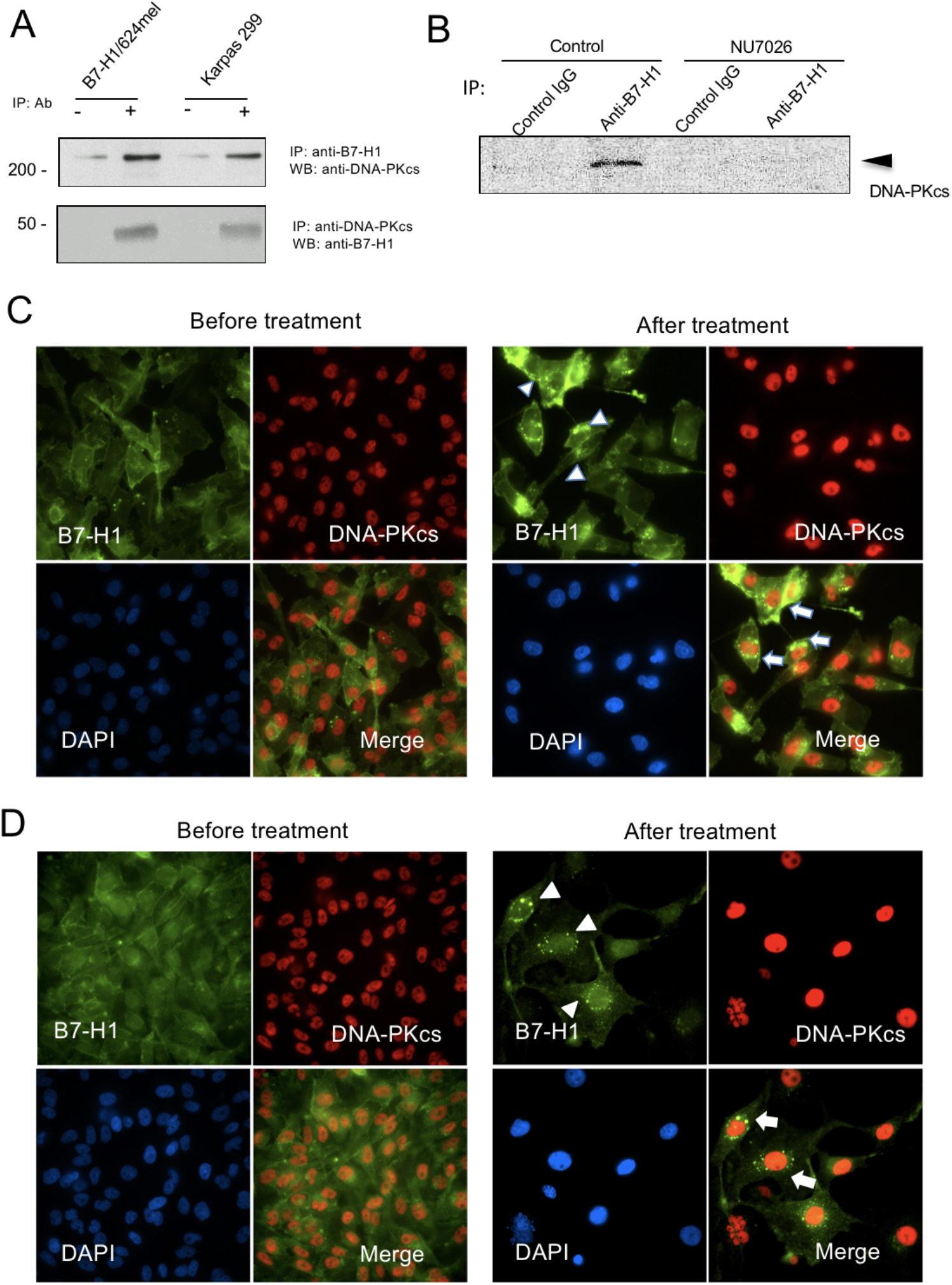
B7-H1 was associated with DNA-PKcs in tumor cells. (A) Immunoprecipitation (IP) and Western blot (WB) identified an association of B7-H1 with DNA-PKcs in B7-H1/624mel cells and Karpas 299 cells that are B7-H1 positive. (B) The association of B7-H1 with DNA-PKcs was identified in MDA-MB-231 cells and this association was abolished by NU7026 (1 μM), an inhibitor of DNA-PKcs. (C-D) Co-localization of B7-H1 and DNA-PKcs in wild type (C) and B7-H1-GFP transfected (D) MDA-MB-231 cells before and after treatment with cisplatin (40 μM) for 72 hours. Co-localizations of B7-H1 and DNA-PKcs were identified in the nuclei. Arrowheads indicated B7-H1 at perinuclear and intranuclear areas.

To characterize the subcellular localization of B7-H1 in association of DNA-PKcs, we stained B7-H1 and DNA-PKcs in tumor cells before and after treatment with chemotherapy considering B7-H1 expression and distribution may change upon treatment (4). DNA-PKcs mainly localized to the nuclei while B7-H1 (endogenous or transfected) was mainly localized in the cytoplasm (arrows) of MDA-MB-231 cells before drug treatment (Fig. 4C-D). Upon cisplatin treatment, we observed B7-H1 translocations at the perinuclear and intranuclear areas in MDA-MB-231 cells (Fig. 4C-D, arrow heads). B7-H1 was enriched in the nuclei and partially colocalized with DNA-PKcs (Fig. 4C-D, arrows). Similar to a previous report (4), we also observed that doxorubicin induced B7-H1 translocation in MDA-MB-231 cells, and the nuclear colocalization of B7-H1 and DNA-PKcs was much reduced upon treatment with DNA-PKcs inhibitor NU7026 (SF. 3A-B). However, B7-H1 KO MDA-MB-231 cells only demonstrated modest increased sensitivity to doxorubicin treatment (SF. 3C), suggesting the impact of B7-H1 translocation on chemoresistance could be drug-specific.

Since DNA-PKcs can activate MAPK/ERK pro-survival signaling pathways(15, 16), the association of B7-H1 and DNA-PKcs prompted us to examine whether overexpression of B7-H1 alters MAPK/ERK signaling pathway in our systems. First, we screened the relative levels of phosphorylated proteins involved in the MAPK/ERK pathway in B7-H1/624mel cells and Mock/ 624mel cells using antibody arrays (R&D Systems). Among 26 proteins, phospho-ERK1/2, phospho-RSK1/2 (downstream targets of ERK pathway), phospho-p38 MAPK and phospho-mTOR were significantly increased among B7-H1/624mel cells compared with Mock/ 624mel cells (Fig. 5A). Given that PI3K/ mTOR pathway has been studied previously by others (17), here we shall focused on our new findings on ERK and p38MAPK.

**Figure 5.**
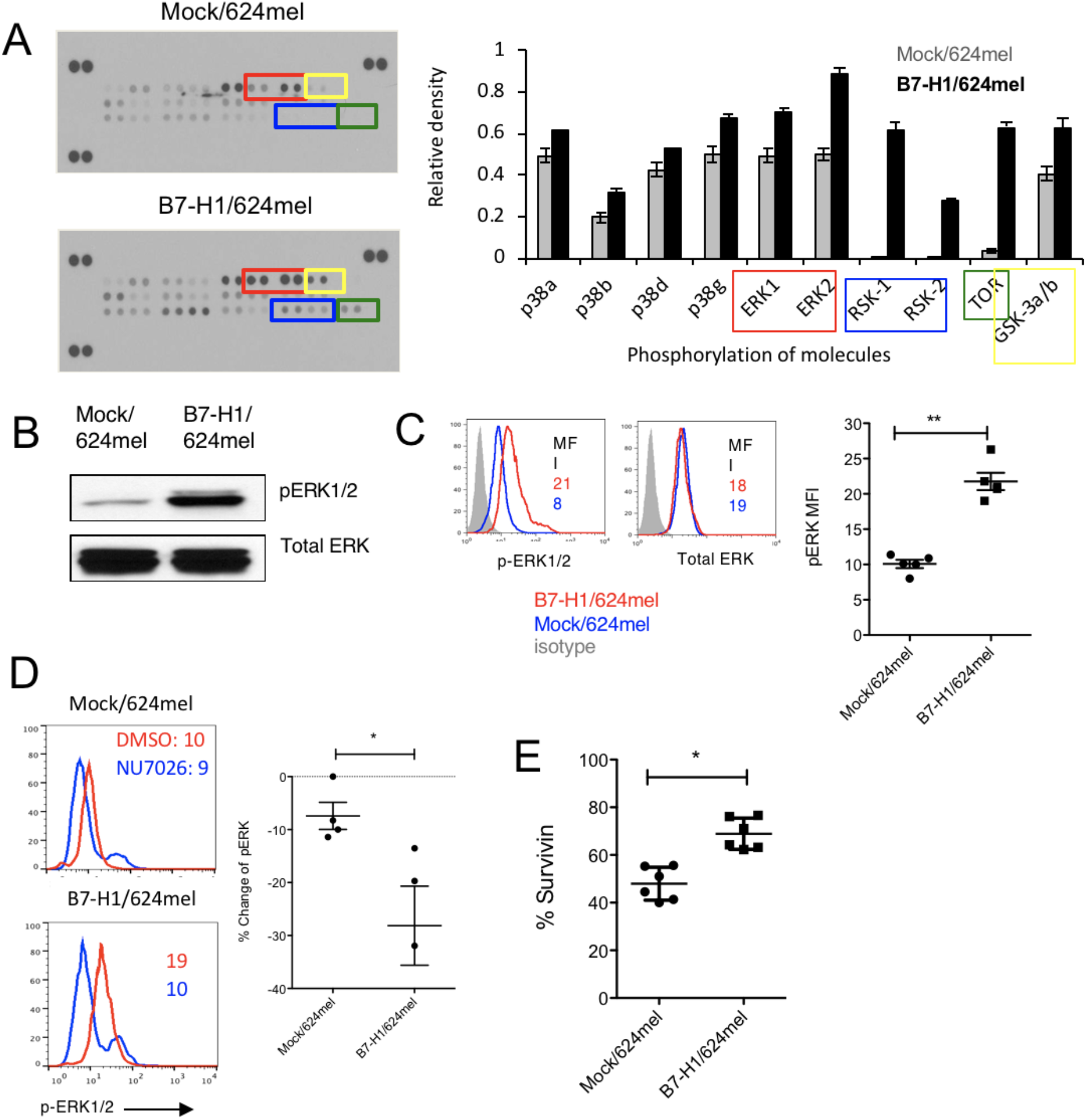
Overexpression of B7-H1 enhanced the activation of ERK pathway. (A) Antibody array assay of phosphorylation of MAPK/ERK pathway. The relative phosphorylation levels of proteins involved in the MAPK/ERK pathway in B7-H1/624mel cells and Mock/ 624mel cells using antibody arrays (R&D Systems). (B-C) The levels of phospho-ERK and total ERK were analyzed by Western blotting (B) and flow cytometry (C). Numbers are mean fluorescence intensity (MfI). **P<0.01 compared between Mock and B7-H1/624mel cells. (D) Histograms show the expression of phospho-ERK after incubation with NU7026 (10 μM, 24 hours). *P<0.05 compared between % changes of phosphor-ERK in Mock and B7-H1/624mel cells. (E) Survivin expression increased in B7-H1/624mel cells in comparison with Mock/624mel (*P<0.05).

The increase of ERK1/2 activation was confirmed by Western blotting assay and intracellular flow cytometry assay that showed a >2-fold increase of phospho-ERK1/2 (Fig. 5B-C). In both flow cytometry and Western blot assays, the total ERK levels remained comparable between B7-H1/624mel and Mock/ 624mel cells. To examine the activity of DNA-PKcs on ERK activation, we treated tumor cells with a DNA-PKcs specific inhibitor (NU7026). The results of Fig. 5D show that NU7026 dramatically decreased (~2-fold) the activation of ERK1/2 in B7-H1/ 624mel cells but not in Mock/ 624mel cells. To examine the consequences of increased activation of ERK, we measured the expression of survivin, an anti-apoptotic molecule and a downstream target of ERK1/2 (18). We found that the expression of survivin was significantly increased in B7-H1/624mel compare to Mock/ 624mel cells (Fig. 5E). Our results suggest at least in part that B7-H1 mediated ERK1/2 activation resulting in enhanced anti-apoptotic programs hence chemoresistance in those cancer cells

We also examined the ERK/MAPK pathway in B7-H1 deficient tumor cells in association with DNA-PKcs. In contrast to 624mel melanoma cells with B7-H1 overexpression, B7-H1 deficient MDA-MB-231 breast cancer cells expressed comparable levels of phospho-ERK as B7-H1 wild type cells, but expressed lower levels of phospho-p38 MAPK (Fig. 6A). The quantitative levels of phospho-ERK and phospho-p38 MAPK were confirmed with flow cytometry assays (Fig. 6B) that also demonstrated less activation of p38 MAPK in B7-H1 deficient tumor cells. To examine whether the activation of p38 MAPK is dependent on the association of B7-H1 and DNA-PKcs, we measured the phospho-p38 MAPK in the presence of NU7026 that can disrupt the association of B7-H1 and DNA-PKcs (Fig. 6B,D). Interestingly, Inhibitor Nu7026 significantly reduced the activation of p38 MAPK in wild type MDA-MB-231 cells but not in B7-H1 deficient cells (Fig. 6C). We observed that the consequences of decreased activation of p38 MAPK led to reduced expression of Bcl-2, a pro-survival molecule in B7-H1 deficient MDA-MB-231 cells compared to wild type cells (Fig. 6D). The decreased levels of Bcl-2 may explain why B7-H1 deficient MDA-MB-231 cells were more susceptible to cisplatin-induced apoptosis (Fig. 6E-F). Taken together, our results suggest that B7-H1-mediated chemoresistance is cell-context dependent and requires ERK/MAPK pathway through the association of B7-H1 with DNA-PKcs (Fig. 7).

**Figure 6.**
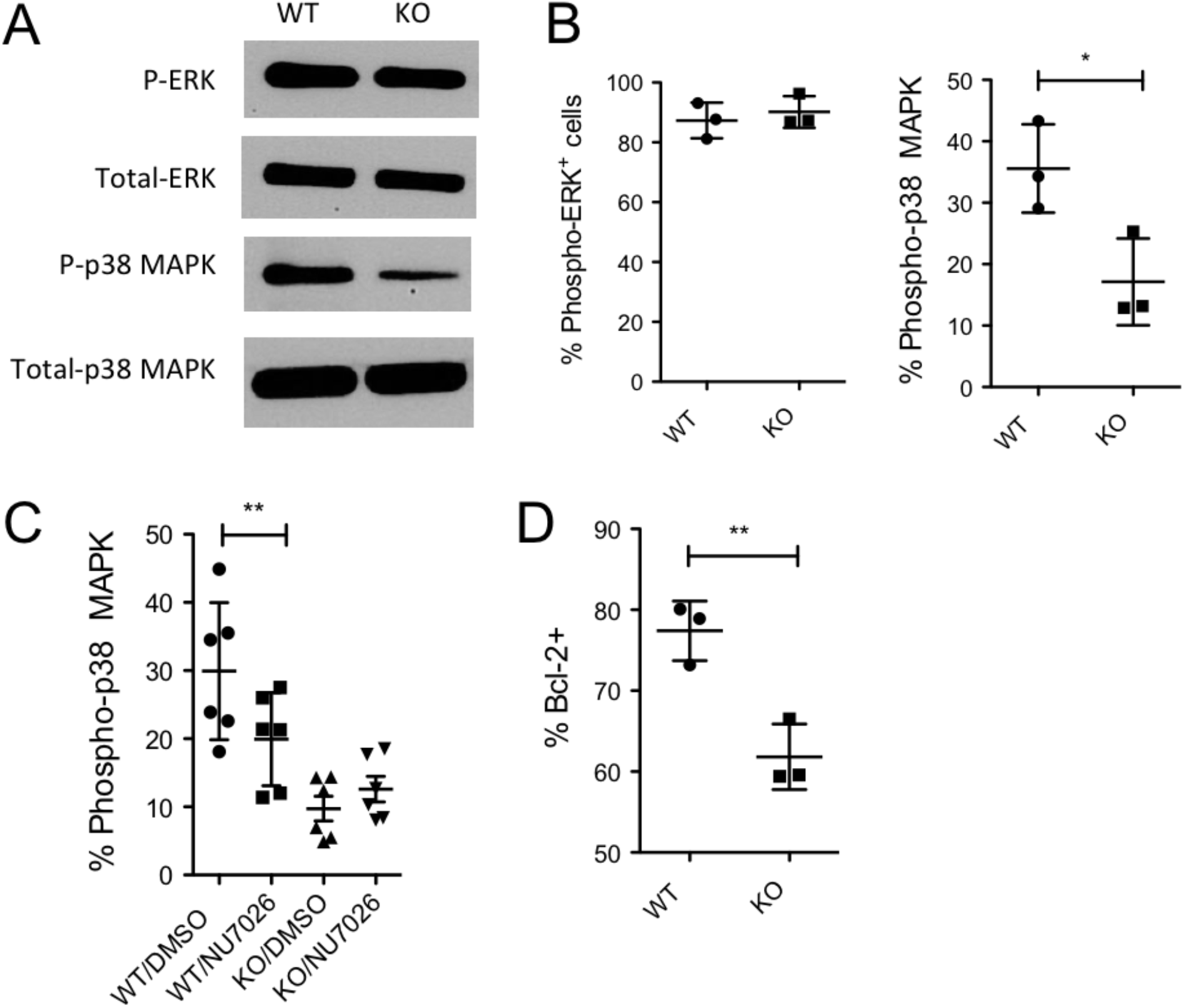
B7-H1 deficiency reduced the activation of p38 MAPK. (A-B) The levels of phospho-ERK or phosphor-p38 MAPK and total ERK or total p38 MAPK in WT or B7-H1 deficient MDA_MB-231 cells were analyzed by Western blotting (A) and flow cytometry (B). *P<0.01 compared between B7-H1 WT and KO MDA-MB-231 cells. (C) The expression of phospho-p38 after incubation with NU7026 (10 μM, 24 hours). **P<0.01 compared between NU7026 treated or not treated WT MDA-MB-231 cells. (D) Bcl-2 expression decreased in B7-H1 KO MDA-MB-231 cells compared to WT cells (**P<0.01).

**Figure 7.**
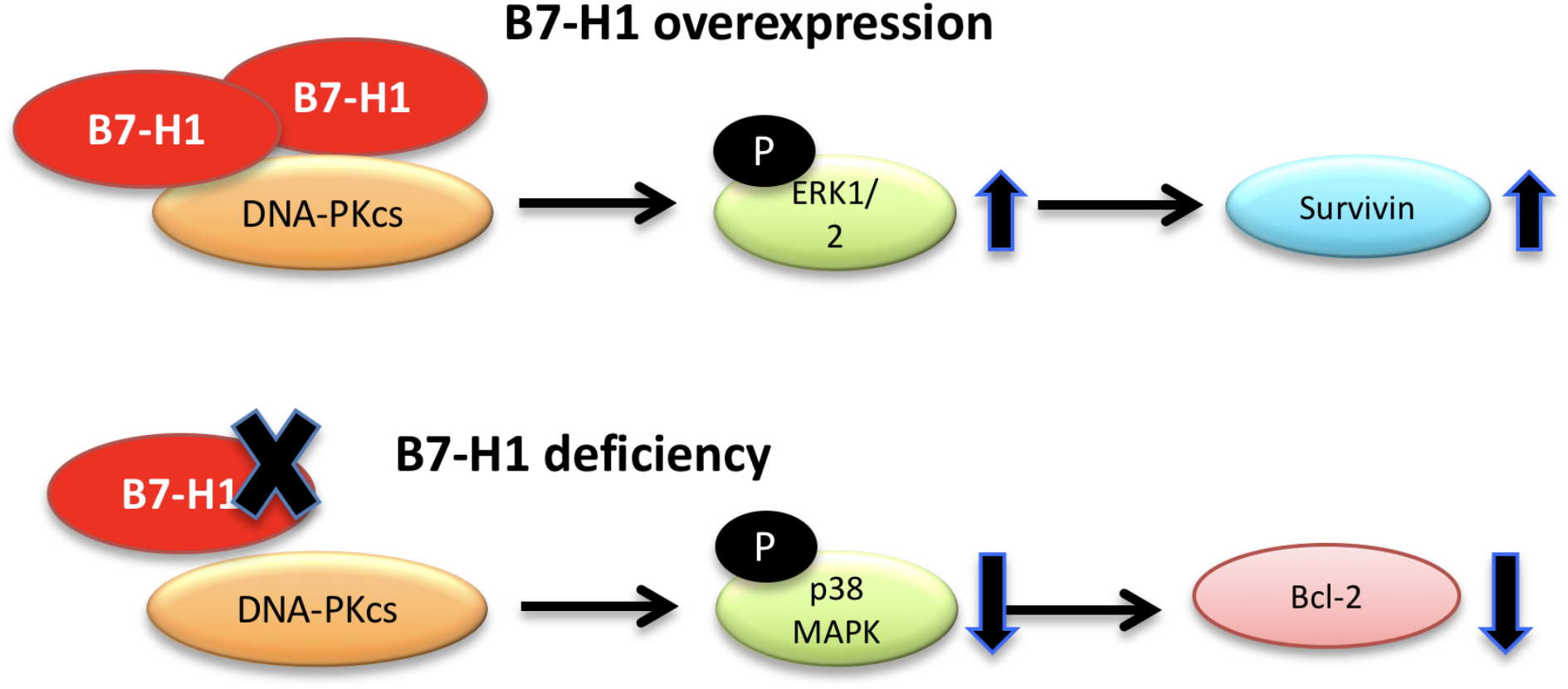
Diagram of potential mechanisms of action of B7-H1 in tumor cell survival and chemoresistance. Overexpression of B7-H1 increases ERK activation that is dependent on DNA-PKcs activity and leads to increased expression of survivin. The deficiency of B7-H1 decreased the activation of p38 MAPK that is dependent on the association of DNA-PKcs and B7-H1 and leads to downregulation of Bcl-2. In both situations, B7-H1 contributes to chemoresistance by promoting or maintaining tumor cell survival through MAPK/ERK pathway.

## DISCUSSION

In this report we identified a new mechanism of B7-H1-mediatd chemoresistance in human tumor cells. We found that overexpression of B7-H1 resulted in increased activation of the ERK pathway in melanoma cells and that steady state expression of B7-H1 is required for activation of p38 MAPK. Regulation of the ERK/MAPK pathway by B7-H1 seems to be dependent on the association of B7-H1 with DNA-PKcs, since pharmacological inhibition of DNA-PKcs readily reduces the activation of ERK or p38 MAPK in wild type tumor cells but not in B7-H1 negative or deficient tumor cells. Thus, our results provided new insights to B7-H1-mediate chemoresistance in cancer cells and suggest that targeting B7-H1 with monoclonal antibody may improve the efficacy of cancer chemotherapy in B7-H1 positive cancer.

Our results are supportive of a previous report that implied the MAPK/ERK pathway might be a target of B7-H1 mediated modulation in tumor cells(19). As ERK activation plays key roles in cell survival and drug resistance(20, 21), the increased ERK activation in B7-H1 overexpressing tumor cells suggests that B7-H1 may promote the ERK pathway in tumor cells to survive upon cytotoxic chemotherapy. In line with B7-H1 overexpressing tumor cells (B7-H1/ 624mel), tumor cells (MDA-MB-231) that constitutively express high levels of B7-H1 also contain high levels of ERK activation. However, since MDA-MB-231 cells harbor mutant *ras* that leads to a high activation of ERK(22), B7-H1 deficiency alone was not able to reduce the activation of ERK in MDA-MB-231. In that regards, our results suggest that B7-H1 may increase ERK activation in tumor cells that contain low degree of ERK activation (like in 624mel cells), but is not required in maintaining ERK activation in tumor cells where ERK activation is high (like in MDA-MB-231 cells).

It has been reported that B7-H1 has a function beyond the negative regulation of T cell responses through its receptor PD-1 (23). Ligation of B7-H1 by antibody causes B7-H1 internalization and disrupts the role of B7-H1 in maintenance of Akt/mTOR signaling (23). To that end, Kleffel et al identified that expression of PD-1 in melanoma cells modulates downstream mTOR signaling (24). Clark et al extended the regulatory of B7-H1 to autophagy in tumor cells by showing B7-H1 sensitizes mouse melanoma cells to pharmacologic autophagy inhibitors in vivo (17). In contrast to these studies using PD-1 positive tumor cells, we addressed the B7-H1 downstream signaling in the absence of PD-1 or CD80, two receptors of B7-H1. We found a new partner of B7-H1 within tumor cells, i.e. DNA-PKcs, which may link B7-H1 with ERK/MAPK pathway. Overexpression of B7-H1 resulted in increased activation of ERK in tumor cells, and B7-H1 deficiency reduced p38 MAPK activation. In addition, DNA-PKcs inhibitor suppressed the activation of ERK and p38 MAPK in the presence of B7-H1 in tumor cells. Since DNA-PKcs or p38 MAPK inhibitors did not change the expression of B7-H1 in tumor cells (supplemental Figure. Xx), it seems that both of DNA-PKcs and p38 MAPK are at the downstream of B7-H1 pathway in tumor cells.

DNA-PKcs is localized to the nucleus where it regulates DNA damage repair in response to cytotoxic drugs(25). However, functions of DNA-PKcs in regulation of pro-survival signaling pathways have been reported in activation of ERK and AKT pathways either in the nucleus or at the cytoplasm(15, 16). Our studies suggest that both ERK and p38 MAPK activation seems function as downstream signal molecules as inhibition of DNA-PKcs activity abolished their activation in the presence of B7-H1. The translocation of B7-H1 induced by chemotherapy in the nucleus of tumor cells suggest that B7-H1 is used as tumor-intrinsic signal molecule to transmit signals to nucleus to prepare risk management at genetic levels. Since B7-H1 confers tumor cell resistance to chemotherapeutic agents that cause DNA damage (cisplatin), the association of B7-H1 with DNA-PKcs may have implication for other mechanisms in B7-H1-mediated chemoresistance beyond the ERK/MAPK pathway.

Of note, B7-H1-mediated chemoresistance is cell-context dependent based on the drug selection. We found that knocking out B7-H1 sensitizes tumor cells (MDA-MB-231 and 786-0) with high steady state expression of B7-H1 protein became more susceptible to cisplatin more so than tumor cells (A549) with low expression B7-H1. These results suggest that high expression of B7-H1 may build a scaffold to maintain high activation of p38 MAPK pathway that contributes to retain high level of Bcl-2(26), an anti-apoptosis molecule. Disruption of B7-H1 expression in these cells would render them more susceptible to chemotherapy as we showed that anti-B7-H1 antibody (H1A) increased the sensitivity of triple negative breast cancer cells to cisplatin in vivo. However, in tumor cells with low or non-B7-H1 expression, disruption of B7-H1 may not significantly affect the sensitivity to cisplatin due to lack of baseline B7-H1-dependent anti-apoptosis signal pathway. In mouse breast cancer cells, B7-H1 deficiency did not increase their susceptibility to cisplatin but did to paclitaxel in vitro and in vivo. The differences between human and mouse breast cancer cells are cancer type (triple negative cancer in human and Her2 positive in mouse) and base levels of B7-H1 expression (higher B7-H1 expression in human than in mouse cells). Our studies indicate that clinical optimization of chemotherapy drugs should consider B7-H1 expression by cancer cells in order to overcome B7-H1-mediated chemoresistance.

In summary, our studies identify a new mechanism of B7-H1-mediated chemoresistance in cancer cells. Both ERK and p38 MAPK pathways together with previously described AKT / mTOR pathway are exploited by either overexpression or steady state expression of B7-H1 to provide anti-apoptosis signals for tumor cells and render tumor cells resistant to chemotherapy. The clinical implication of our study is that the status of B7-H1 expression by cancer cells should be considered in optimizing chemotherapy in order to overcome B7-H1-mediated chemoresistance. Monoclonal antibody to B7-H1/PD-L1 may have the potential to increase the sensitivity of cancer cells to chemotherapy along with its function to restore T cell immunity to cancer.

## METHODS AND MATERIALS

### Cell lines and reagents

Human cancer cell lines (MDA-MB-231, 786-0, A549) were purchased from ATCC (Manassas, VA). Tumor cells were cultured and maintained in medium indicated by ATCC. B7-H1 or OVA (mock) transfected 624mel cells were cultured in RPMI 1640 medium (Cellgro) and supplemented as described previously (9). Cells were cultured in a 37°C humidified chamber at 5% CO2. Chemotherapy drugs were purchased form Mayo Pharmacy or Sigma.

### B7-H1 transfection and knockout

Human B7-H1 was knocked-out by CRISPR/ Cas9 technology. The guide sequence (5’-ATTTACTGTCACGGTTCCCA-3’) specific to human B7-H1 exon 3 (second coding exon), designed using CRISPR DESIGN tool (http://crispr.mit.edu) and cloned into px458 plasmid coexpressing GFP (Addgene, #52961). Thirty six hours after transfection, cells were sorted for GFP and sub-cloned using flow cytometry. Two weeks later, single cell subclones were genotyped by PCR and validated Western blotting for B7-H1 protein depletion. B7-H1 expression level was determined by flow cytometry and Western blotting.

### Immunofluorescence staining

Following growth on PBS and medium pre-rinsed coverslips, cells were fixed with 4% formalin or paraformaldehyde for 15 min., washed 4x with PBS, and permeabilized for 10 min. with 0.2% Triton X-100 or 0.5% IGEPAL ca-360. After washing with PBS, cells were blocked with 3% milk/PBS, then incubated at 4°C overnight with primary antibodies (1:100 anti-DNA-PKcs and 1:300 anti-B7-H1 antibody 5H1) diluted in blocking solution. Six 3% milk/PBS washes were performed prior to 1 hour incubation with secondary antibody (Life Technologies Fluorescein-conjugated goat anti-mouse and Alexa 594-conjugated goat anti-rabbit IgG) diluted 1:100 in blocking solution. Following five PBS washes, re-fixation for 10 min. with 4% paraformaldehyde, and two dH2O washes, coverslips were mounted with SlowFade Gold antifade reagent with DAPI (Invitrogen) and cured for 24 hrs in dark at RT. Nail-polish sealed coverslips were visualized using a Zeiss LSM 510 confocal microscope.

### MTS cytotoxicity assay

1 × 10^4^ cells were seeded into 96-well plates and chemo-drug was applied. Following 72 hour incubation, 20 μl/ well CellTiter 96 Aqueous One Solution Reagent (Promega) was added. After 2 hours of incubation, absorbance at 490 nm was recorded using an ELISA plate reader. Control and all concentrations of drug were assayed in triplicate, and the absorbance at each drug concentration was normalized relative to that of untreated controls.

### Flow cytometry analysis

Fluorochrome-conjugated Abs against human B7-H1 (MIH1), PD-1 (EH12.2H7) and CD80 (L307.4) were purchased from BD Biosciences (Mountain View, CA), BioLegend (San Diego, CA), or eBioscience (San Diego, CA). To detect intracellular Survivin (clone 32.1, Novus), and Bcl-2 (Clone 50E3, Cell Signaling Inc.), tumor cells were incubated in Fixation Buffer (BioLegend) for 20 min at room temperature, followed by permeabilization using Permeabilization Wash Buffer (BioLegend). After staining with antibody, cells were washed three times with washing buffer before analysis. Apoptosis of tumor cells was analyzed by staining with Annexin V (BD Biosciences) and tetramethylrhodamine (TMRE; ethyl ester) (Invitrogen-Molecular Probes) for 10 min at room temperature. At least 100,000 viable cells were live gated on FACSCailbur (BD Biosciences) instrumentation. Flow cytometry analysis was performed using FlowJo software (Tree Star, Ashland, OR).

### Measurement of phosphorylation of ERK and p38 MAPK

Human Phospho-MAPK Array Kit (R&D Systems) was used to screen phosphorylated proteins between B7-H1 and Mock transfected 624mel cells. After incubation, cells are fixed in 4% formaldehyde for 10 min at 37 °C followed with permeabilization in ice-cold methanol for 30 min on ice. After washing, cells were blocked with Fc receptor blockers and incubated with rabbit mAb to phospho-ERK (T202/Y204) (clone 197G2) or p38 MAPK (Thr180/Tyr182) (Clone D3F9, Cell Signaling #4511) or Isotype control IgG for 1 hour at room temperature, followed with incubation with secondary anti-rabbit IgG (H+L), F(ab’)_2_ Fragment (Alexa Fluor^®^ 488 Conjugate) (Cell Signaling #4412) for 30 min at room temperature. To inhibit the activity of DNA-PKcs, NU7026 (Selleckchem #S2893) was added at 10 μM in cell culture an hour before cell culture. The intracellular levels of ERK or p38 MAPK activation was analyzed by flow cytometry.

### Immunoprecipitation (IP), Western blotting and Mass spectrometry

Immunoprecipitation of endogenous proteins that may be associated with B7-H1 was performed using B7-H1/ 624mel and B7-H1 positive Karpas-299 cells (a human T cell lymphoma cell line) purchased from American Type Culture Collection and propagated in complete media (RPMI, 10% FBS, 20 mmol/L HEPES, and penicillin/ streptomycin). Cells were harvested in 1%NP40 lysis buffer containing 50 mM Tris-HCL pH 8.0, 150 mM NaCl, 5 mM EGTA, 5 mM EDTA, 30 mM MgCl2, 1.3% Beta-glycerophosphate, 1 mM DTT, 0.1 mM Na-Vanadate, 0.1 mM HaF, 0.4% p-nitrophenyl phosphate and protease inhibitors. Extracts were incubated overnight with protein G beads pre-coated with anti-B7-H1 mAb (5H1), control IgG or polyclonal anti-DNA-PKcs antibody (H163, Santa Cruz Biotechnology, Santa Cruz, CA). The proteins eluted from the protein G beads were resolved 5-7.5% SDS gel and detected by *Coomassie Blue* and Western blotting. Rabbit anti-DNA-PKcs antibody at 1:1000 dilution (H-163, Santa Cruz Biotechnology) and mouse anti-B7-H1 antibody at 1:500 dilution (H1A or B11, established at Dong lab) were used as the primary antibodies for Western blotting. Mass spectrometry analysis of protein bands were performed at Mayo Clinic Proteomics Research Core Facility.

### Animal studies

Both B7-H1 WT and KO MDA-MB-231 cells were subcutaneously injected into SCID mice (Jackson lab) at 2×10^6^ cells per mouse. Cisplatin was administrated intraperitoneally (i.p.) at 5 mg/kg for 2 doses on days 7 and 13. Anti-human B7-H1 monoclonal antibody (clone H1A(14), produced at Dong lab) or isotype control (mouse IgG) was i.p. injected at 200 μg/ mouse on days 6, 9, 12, 15 and 18. B7-H1 wild type or knockout tumor cells were isolated from wild type breast cancer NeuT mice provided by Dr. L. Pease (Mayo Clinic, Rochester, MN) or from B7-H1 knockout NeuT mice (Dong lab). Mouse breast tumor cells (5 x10^5^ cells) were injected subcutaneously at the right flank of Balb/c purchased from Taconic Farms (Germantown, NY) and used at age of 12 weeks. On day 8 and 14, paclitaxel (Taxol) was injected i.p. at 25 mg/kg, and control groups were treated with saline. Tumor growth was evaluated every 2-3 days until days 35-40 when all mice were euthanized that were not already sacrificed for rapid tumor growth or ulceration. In compliance with animal care guidelines, mice were euthanatized when either primary or secondary tumors reached ethical endpoints or developed ulcerations. Mayo Clinic’s Institutional Animal Care and Use Committee approved this study.

### Statistical analysis

Mann-Whitney test or unpaired two-side t test was used to compare phenotype between independent groups (treatments) or cell populations (B7-H1 WT and KO cells). The drug cytotoxicity assay was analyzed by the area under curve between the control line and the tested drugs was calculated between B7-H1 WT and KO cells. The therapeutic effects of chemotherapy were analyzed by two-way ANOVA because two independent factors (B7-H1 phenotype or drug treatment) were analyzed. All statistical analyses were performed using GraphPad Prism software 5.0 (GraphPad Software, Inc., San Diego, CA). A *P* value <0.05 was considered statistically significant.

## AUTHOR CONTRIBUTIONS

Conception and design: X. Wu, L. Wang, K. Ling, H. Dong

Dong Development of methodology: X. Wu, Y. Li, L. Wang, H. Dong

Acquisition of data: X. Wu, Y. Li, X. Liu, S. Cao, C. Chen, S. Harrington.

Analysis and interpretation of data: X. Wu, Y. Li, A. Mansfield, S. Park, R. Dronca, E. Kwon, Y. Yan, L. Wang, K. Ling, H. Dong

Writing, review and revision of manuscript: X. Wu, L. Wang, K. Ling, A. Mansfield, S. Park, Y. Yan, R. Dronca, H. Dong

Study supervision: L. Wang, K. Ling, H. Dong

---

**Supplemental figure 1.**
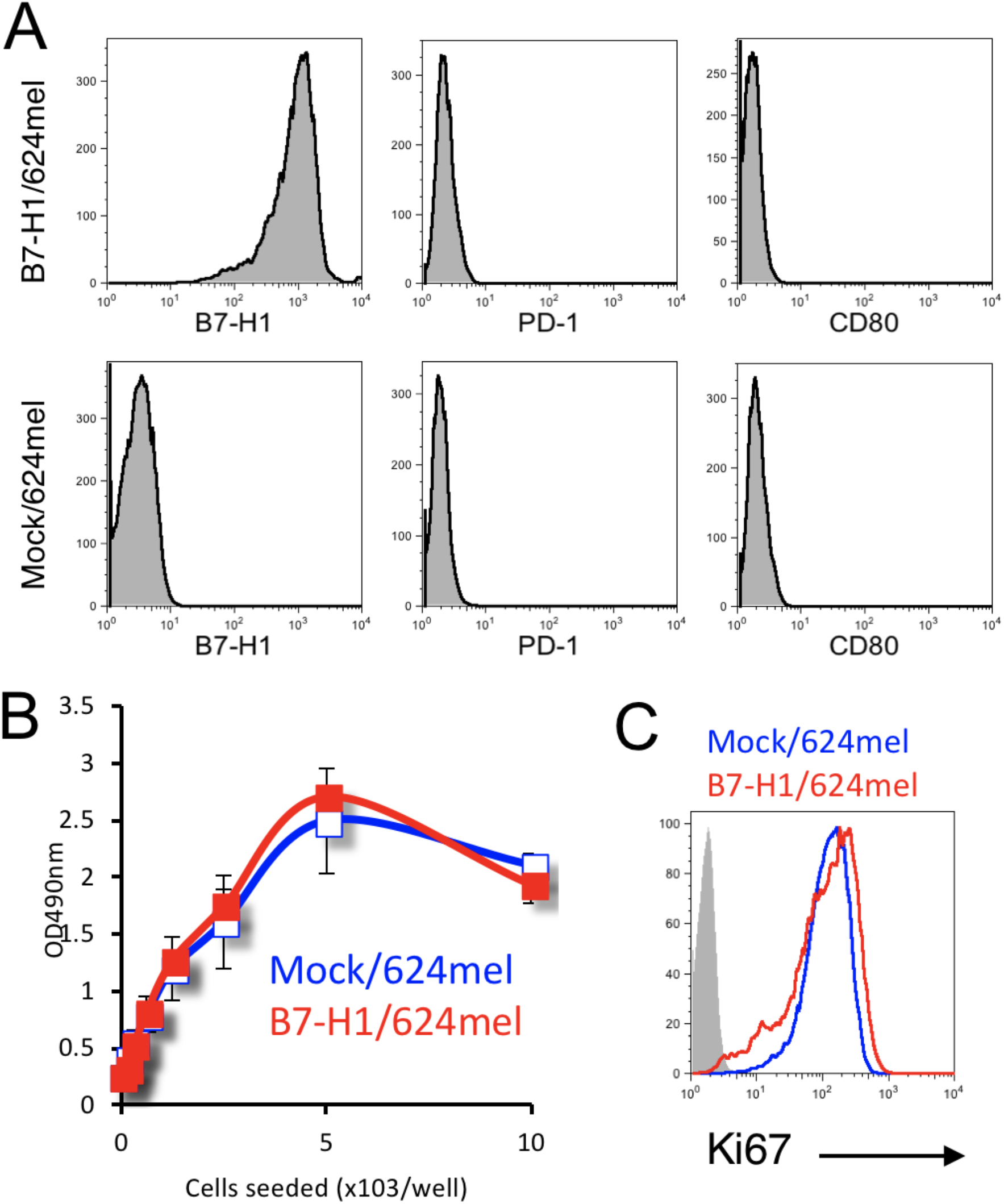
(A) No PD-1 and CD80 (B7-1) expression by Mock/624mel and B7-H1/ 624mel cells. (B-C) Comparable growth rate between B7-H1/624mel and Mock/ 624mel cells in vitro. The proliferation of both tumor cells was analyzed by MTS assay after 72h of culture (B) and flow cytometry assay for the expression of intranuclear protein Ki67 (C).

**Supplemental figure 2.**
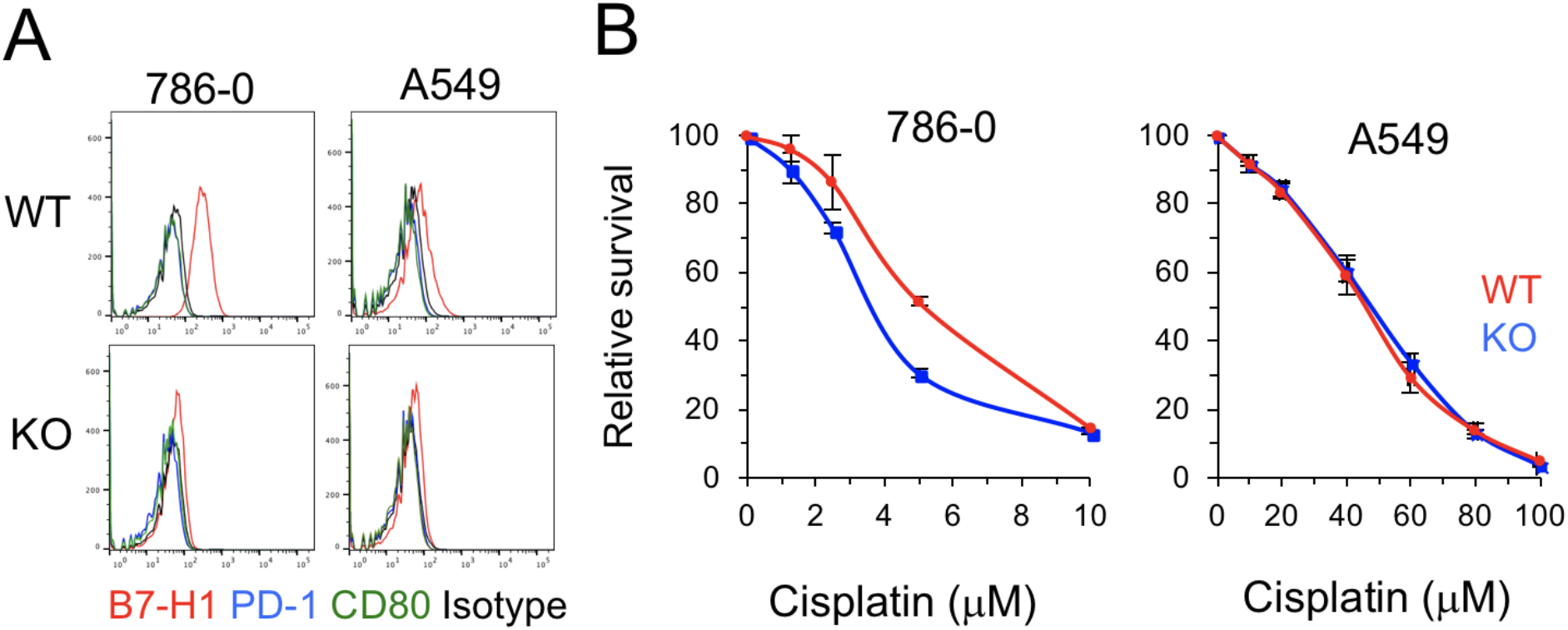
(A) B7-H1, PD-1 and CD80 expression in wild type or B7-H1 KO 786-0 and A549 tumor cells. (B) B7-H1 KO 786-0 cells, but not B7-H1 KO A549 cells, are more sensitive to cisplatin compared to their WT cells as determined by MTS assay. The p-value for area under the curve (dose-response curves) is significant (p<0.01) in treatment with cisplatin in B7-H1 KO 786-0 cells.

**Supplemental figure 3.**
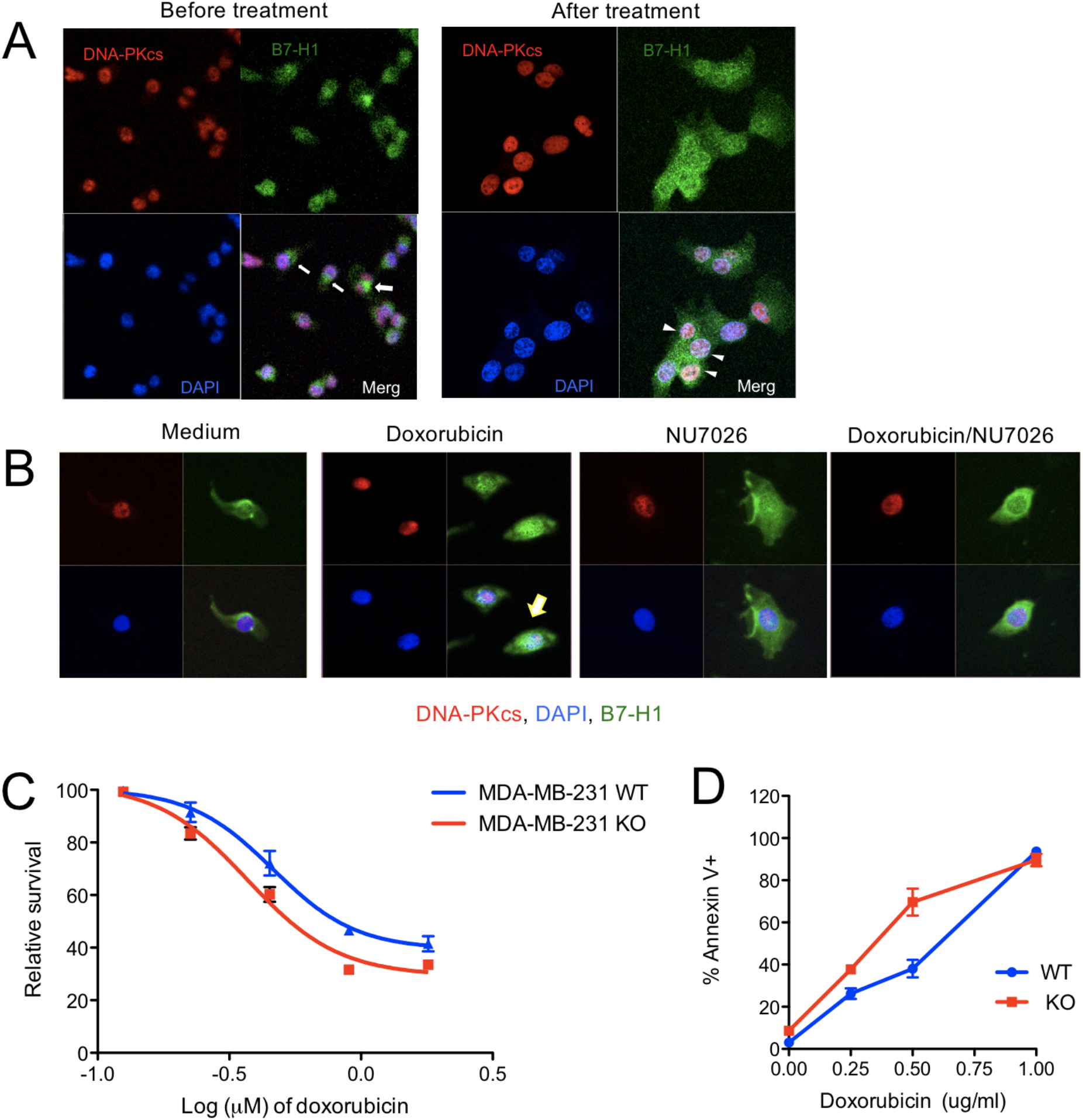
(A) Co-localization of B7-H1 and DNA-PKcs. MDA-MB-231 cells were treated with or without doxorubicin (2 μg/ ml) for 2 hours. Co-localizations of B7-H1 and DNA-PKcs were identified in the nuclei. Arrows or arrowheads indicate B7-H1 in plasma or in nuclei, respectively. (B) DNA-PKcs inhibitor NU7026 disrupted the co-localization of B7-H1 and DNA-PKcs were identified in the nuclei of tumor cells. Arrow indicates B7-H1 in nuclei with DNA-PKcs. (C) B7-H1 deficient MBA-MD-231 cells are modestly sensitive to doxorubicin than WT cells as determined by MTS assay.

